# Dimerisation and twist reversal of the Lewy fold in α-synuclein mutants with Parkinson’s disease and dementia

**DOI:** 10.64898/2026.04.07.716933

**Authors:** Hongwei Zhang, Alexey G. Murzin, Jennifer A. Macdonald, Tiara V. Hinton, Sew Peak-Chew, Catarina Franco, Patrick W. Cullinane, Thomas Warner, Ayami Okuzumi, Raquel Real, Kenya Nishioka, Daisuke Taniguchi, Daita Kaneda, Huw Morris, Henry Houlden, Bernardino Ghetti, Nobutaka Hattori, Zane Jaunmuktane, Yang Yang, Sjors H.W. Scheres, Michel Goedert

**Affiliations:** Department of Structural Biology, Van Andel Institute, Grand Rapids, MI, USA; Medical Research Council Laboratory of Molecular Biology, Cambridge, UK; Department of Clinical and Movement Neurosciences, UCL Queen Square Institute of Neurology, London, UK; Queen Square Brain Bank for Neurological Disorders, UCL Queen Square Institute of Neurology, London, UK; Department of Neurology, Juntendo University Faculty of Medicine, Tokyo, Japan; UCL Movement Disorders Centre, University College, London, UK; Department of Radiology, National Centre for Geriatrics and Gerontology, Aichi, Japan; Department of Neuromuscular Diseases, UCL Queen Square Institute of Neurology, London, UK; Department of Pathology and Laboratory Medicine, Indiana University School of Medicine, Indianapolis, IN, USA

## Abstract

Dominantly inherited missense and gene dosage mutations in *SNCA*, the α-synuclein gene, cause familial forms of Parkinson’s disease and dementia. Here we report the structures of α-synuclein filaments from the brains of such individuals. Pathogenic mutations A53T and G51D in *SNCA* give rise to singlets and doublets of the Lewy fold with a left-handed helical twist in the absence of a peptide-like density for island A. By contrast, filaments from the non-pathogenic variant H50Q consist of singlets of the right-handed Lewy fold with a density for island A, like filaments of wild-type α-synuclein. The structures of filaments from homozygous mice transgenic for human mutant A53T α-synuclein (line M83) are unlike those from human brains. They are more similar to the multiple system atrophy folds than to the Lewy fold of Parkinson’s disease, Parkinson’s disease dementia and dementia with Lewy bodies

## Introduction

Parkinson’s disease (PD), Parkinson’s disease dementia (PDD), dementia with Lewy bodies (DLB) and multiple system atrophy (MSA) are the major synucleinopathies [1]. They have the presence of abundant filamentous inclusions of α-synuclein in common. Dominantly inherited missense mutations in *SNCA*, the α-synuclein gene, cause PD with and without dementia [2-10]. The same is also true for gene dosage mutations of *SNCA* (duplications and triplications) [11-13]. The existence of missense mutations and multiplications of *SNCA* that cause diseases with abundant filamentous inclusions of α-synuclein has led to the notion that α-synuclein is a key player in all cases of PD, PDD and DLB. In support, genome-wide association studies have shown that single nucleotide polymorphisms in *SNCA* are associated with elevated levels of α-synuclein, which increases the risk of disease [14-16].

A53T in *SNCA* was the first mutation known to cause PDD and is also the most common, with more than twenty families having been described [2,17,18]. G51D in *SNCA* has been identified in four families with PDD [6,7,19,20]. Some clinicopathological characteristics overlapped with MSA. Variant H50Q in *SNCA* has been reported in PD [5,21]. However, unlike cases with the A53T and G51D mutations, the family history has been inconsistent. Moreover, the heterozygous variant in *SNCA* that encodes H50Q α-synuclein has also been found in population databases and was not enriched in PD, questioning its pathogenicity [22].

By electron cryo-microscopy (cryo-EM), α-synuclein filaments from PD with or without dementia shared the Lewy fold [23]. It was made of a single protofilament with a right-handed helical twist that extended from residues 31-100 of α-synuclein. Two proteinaceous densities of unknown amino acid sequence that are not connected to the filament core (islands A and B) were also present. Structural studies on the *in vitro* helical assembly of recombinant α-synuclein [24-28] have suggested that filaments from human brains with pathogenic *SNCA* mutations will adopt folds different from those of assembled wild-type α-synuclein.

Here we show that this is not the case. Instead, we demonstrate a correlation between PD-causing mutations in *SNCA* and dimerisation of the Lewy fold with a left-handed helical twist, in the absence of island A. We also report that the cryo-EM structures of filaments from the brains of homozygous mice transgenic for human A53T α-synuclein (line M83) [29] differ from those of filaments extracted from human brains. They are related more closely to filaments of MSA [30-33] and juvenile-onset synucleinopathy (JOS) [34] than to filaments of PD [23].

## Results

### *SNCA* mutation A53T: Left-handed singlets and doublets of the Lewy fold without island A

We determined the cryo-EM structures of α-synuclein filaments from the temporal cortex of a previously described male with a family history of PDD caused by a heterozygous mutation in SNCA encoding A53T α-synuclein [35] (Fig.1, fig. S1 and table S1). This individual developed symptoms of PD aged 46 and died aged 52. The A53T mutation was confirmed by whole-genome sequencing.

**Fig. 1:**
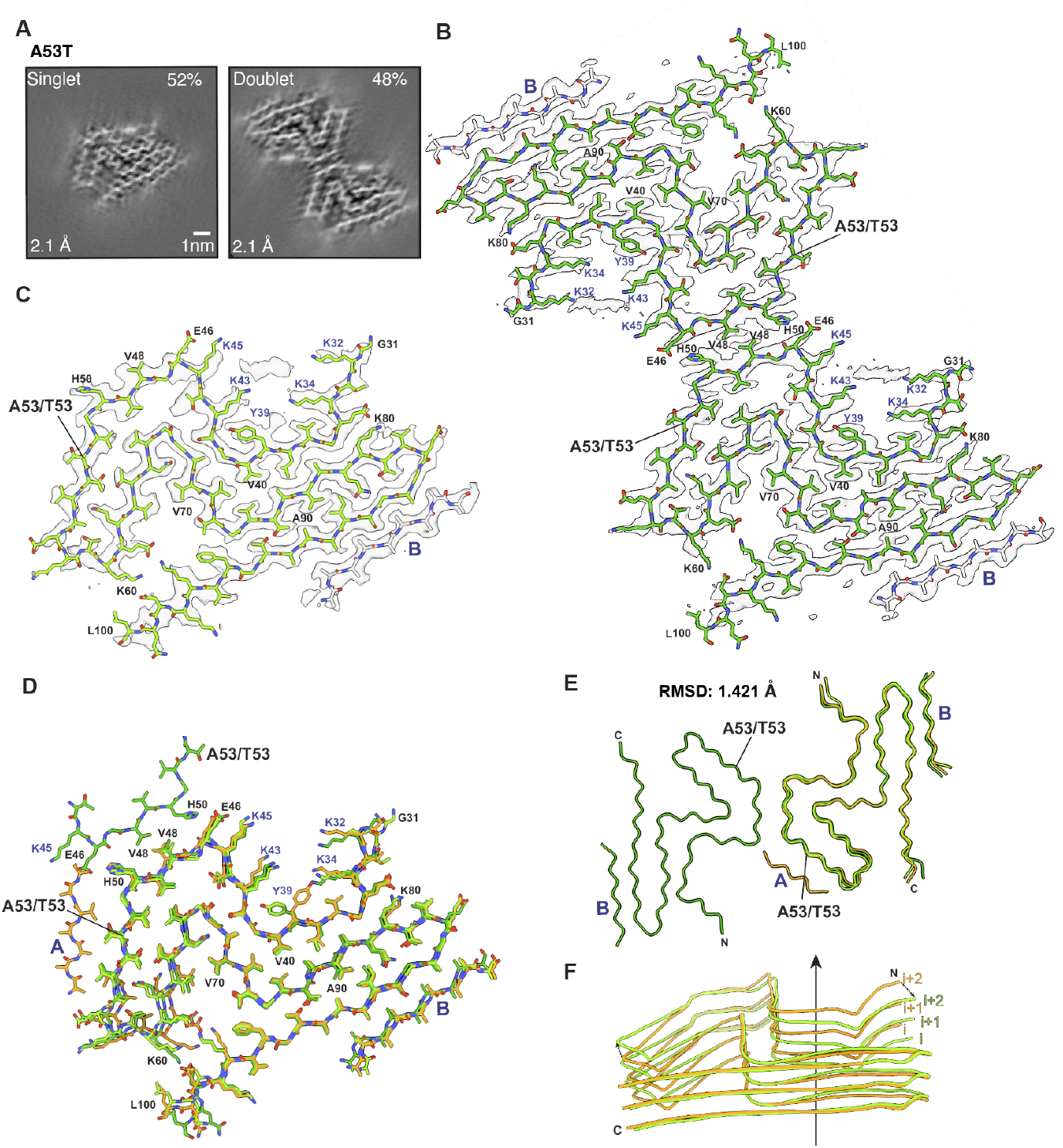
Cryo-EM structures of A53T α-synuclein filaments from human brain. **(A)** Cross-sections through the cryo-EM reconstructions, perpendicular to the helical axis and with a projected thickness of approximately one rung, are shown for α-synuclein singlet and doublet filaments from a case with *SNCA* mutation A53T. Scale bar, 1 nm. **(B)** Cryo-EM density map (transparent grey) and atomic model of the doublet filaments. The model for α-synuclein is shown in dark green and that of island B in grey. **(C)** As in B, but for the singlet, with the atomic model for α-synuclein in light green. **(D)** All-atom superposition of the A53T singlet (light green), part of A53T doublet with its dimer interface (dark green) and the Lewy fold from idiopathic PD, PDD and DLB (orange). **(E)** Top view of an overlay of backbones of the Lewy fold (orange), the A53T singlet (light green) and the A53T doublet (dark green). **(F)** Side views at an angle of 30° from the x tilt axis of the backbones of the Lewy fold (orange) and the A53T singlet (light green), to show differences in Z-heights. The solid arrow indicates the Z-axis, and the dashed arrows indicate the shifts at N-terminus and the hairpin region (residues 51-67).

By 2D class averaging, 49% of filaments lacked the helical twist required for structure determination. As observed for idiopathic PD, PDD and DLB [23], untwisted 2D class averages were consistent with projections of the same structure as the twisted filaments. Cryo-EM structures of the twisted filaments were determined to a resolution of 2.1 Å. Singlet and doublet filaments with diameters of ∼10 nm and ∼20 nm and long crossover distances (∼5,200 Å for singlets and ∼5,500 Å for doublets) were present in roughly equal numbers. Unexpectedly, they adopted a left-handed helical twist, in contrast to the right-handed twist previously described in the Lewy fold singlets from sporadic cases of PD, PDD and DLB [23]. Despite this change in handedness, the backbone conformation of the A53T singlets was like that of the Lewy fold. The two folds superimposed with a backbone root mean square deviation (RMSD) of 1.421 Å (Fig. 1D and E); the (XY) amyloid packing difference (APD) [36] was 4% (Fig. S2A). This small difference arose from a reorientation of the side chain of tyrosine 39. The N-terminal part of the fold, which interacted with an unidentified cofactor density through lysines 32, 34, 43 and 45, rotated ∼8° relative to the helical Z-axis. Differences in the relative Z-coordinates were also observed for the hairpin that was formed by residues 51-67. The doublet A53T filaments comprised two identical protofilaments with the same fold as the singlet that were related by a 2-start (or pseudo-2_1_) helical symmetry (Figure 1B and D), each with a left-handed helical twist. The protofilament interface was formed by residues E46, V48 and H50 from each protofilament. A53T protofilaments contained a non-proteinaceous density in the same position as in the Lewy fold.

Unlike the Lewy fold [23], which contains two disconnected peptide-like densities (islands A and B), the doublet A53T protofilaments lacked the density for island A, thus exposing the mutation site to the solvent. In singlet A53T protofilaments, weak densities were present in the region of island A, but they were not connected in a peptide-like manner. The cryo-EM map of the singlet structure was consistent with the presence of both residues A53 and T53 within filaments, which was confirmed by mass spectrometric analysis of the sarkosyl-insoluble fractions (Fig. S3). A threonine residue at position 53 instead of an alanine replaces a small, nonpolar amino acid by one with a larger side chain containing a hydroxyl group, thus changing the steric and hydrogen-bonding environment. This change may perturb interactions with residues in island A (Fig. 1).

### *SNCA* mutation G51D: Left-handed doublets of the Lewy fold without island A

We next determined the cryo-EM structures of α-synuclein filaments from the frontal cortex of two previously described individuals with a family history of PDD caused by a heterozygous mutation in *SNCA* encoding G51D α-synuclein [20] (Fig. 2, fig. S4 and table S1). Case 1 (family two, patient III) was diagnosed with PD aged 69 and developed cognitive impairment aged 71. This female patient died aged 75. Case 2 (family two, patient IV) was the son of case 1. He was diagnosed with PD aged 46, developed cognitive impairment aged 50 and died aged 52. The G51D mutation was confirmed by whole-genome sequencing.

**Fig. 2:**
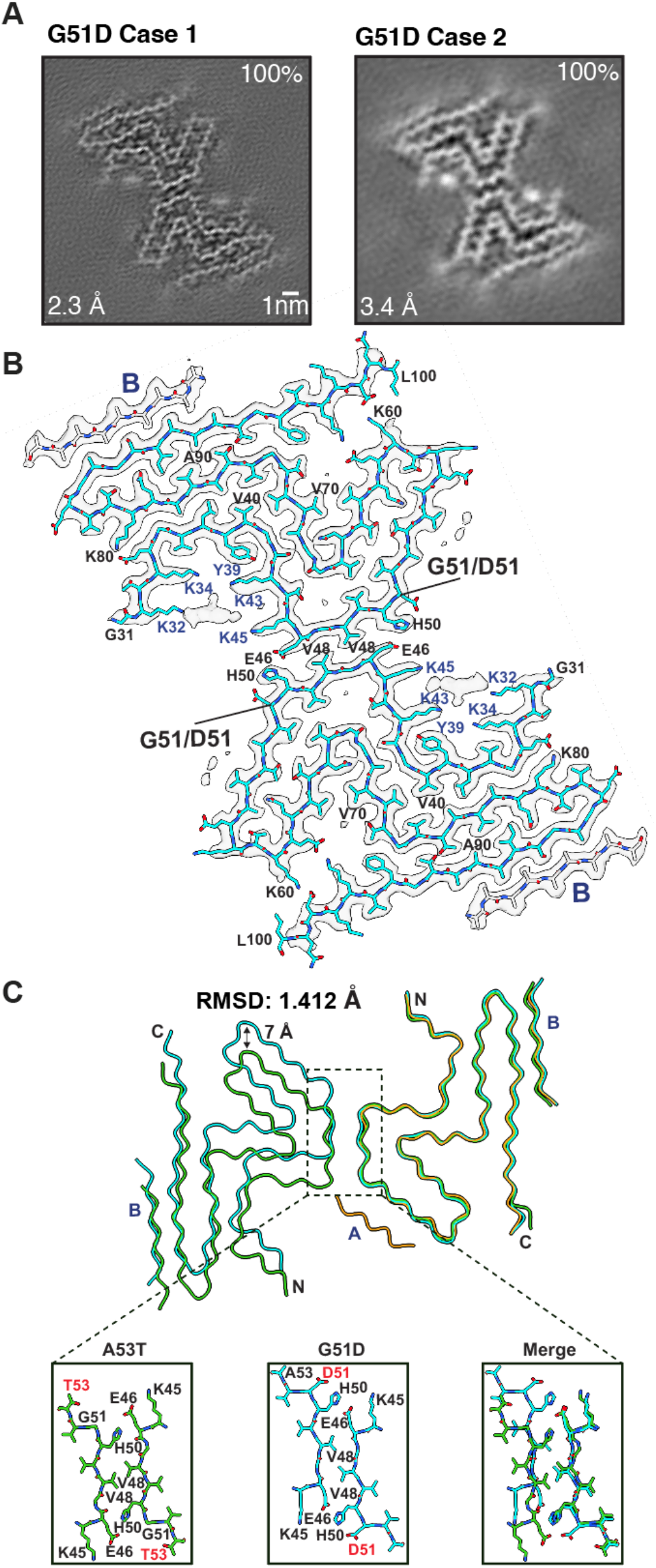
Cryo-EM structures of G51D α-synuclein filaments from human brain. **(A)** Cross-sections through the cryo-EM reconstructions, perpendicular to the helical axis and with a projected thickness of approximately one rung, are shown for α-synuclein doublet filaments from cases 1 and 2 with *SNCA* mutation G51D. Scale bar, 1 nm. **(B)** Cryo-EM density map (transparent grey) and atomic model (cyan) of the doublet filaments of case 1. The model for island B in shown in grey. **(C)** Comparison of the backbone trace of the G51D doublet (cyan), the A53T doublet (green) and the sporadic PD Lewy fold singlet (orange). The zoomed-in views highlight the differences at the dimer interface of the G51D and A53T doublets.

By 2D class averaging, 34% of G51D α-synuclein filaments of cases 1 and 2 were untwisted. Structure determination of the twisting filaments yielded identical doublet cryo-EM structures, with resolutions of 2.3 Å and 3.4 Å. No singlet filaments were observed. The G51D filaments had a left-handed helical twist. They consisted of two identical protofilaments with the Lewy fold that were related by a 2-start helical symmetry. The protofilaments of A53T and G51D superimposed with an RMSD of 0.675 Å. As in the A53T doublet, the protofilament interface was formed by residues E46, V48 and H50 from each protofilament. However, the protofilaments were shifted by 7 Å relative to the A53T protofilaments. G51D dimers contained non-proteinaceous densities in the same position as in the Lewy fold. The G51D protofilament fold also had an APD of 4% with the Lewy fold (Fig. S2B), with similar differences in the orientation of the side chain of tyrosine 39 and differences in the Z-coordinates of the N-terminal part of the fold; the two folds superimposed with an RMSD of 1.412 Å.

Island A was again missing from the dimeric protofilaments, indicative of the structural impact of the G51D mutation. In addition, this mutation probably accounts for the shift of the protofilament interface (Fig. 2C), compared to the A53T doublet filaments. In the G51D interface, the side chain of H50 was more solvent-exposed than in the A53T interface, presumably because it carried a positive charge at neutral pH, stabilised by the negatively charged side chain of the neighbouring D51.

Since the densities of several carboxyl groups were missing from the cryo-EM reconstruction, the presence of residue D51 could not be decided from local density alone. There was a short stump of density, probably corresponding to a Cβ atom, that was smaller than the side chain densities of A53 and D98, the only other aspartic acid residue in the Lewy fold core. This is consistent with both G51 and D51 being present in the filaments, which was confirmed by mass spectrometric analysis of the sarkosyl-insoluble fractions (Fig. S3).

### *SNCA* variant H50Q: Right-handed singlets of the Lewy fold with island A

We also determined the cryo-EM structure of α-synuclein filaments extracted from the amygdala of a previously undescribed case of PD without a family history, but with a heterozygous variant in *SNCA* encoding H50Q α-synuclein (Fig. 3, fig. S5 and table S1). This female developed PD aged 47 and died aged 78. The H50Q variant was identified using the Neurobooster array and confirmed by whole-genome sequencing. The clinical phenotype was consistent with idiopathic PD, with slow disease progression, a sustained levodopa response and no cognitive impairment. Neuropathology also closely resembled that of idiopathic PD, with abundant Lewy bodies and Lewy neurites that were Gallyas-Braak silver-negative (Fig. S6A-D and G). By contrast, the neuronal inclusions from the A53T case and case 2 of G51D were silver-positive (Fig. S6E and F). A case of MSA served as a positive control (Fig. S6H). Small numbers of amyloid-β plaques and tau tangles were also present in the H50Q case. By cryo-EM 2D class averaging, 75% of α-synuclein filaments extracted from the amygdala of the H50Q case were untwisted. The cryo-EM structure of the twisted filaments of α-synuclein was determined to a resolution of 2.9 Å with an APD of 0% (Fig. S2C) and a backbone RMSD of 0.271 Å; it was nearly identical to the wildtype Lewy fold, including islands A and B and the cofactor densities. Like the wildtype α-synuclein filaments [23], the H50Q filaments exhibited a right-handed helical twist. One difference between the two structures was the side chain density at the mutation site (Fig. 3C). In the H50Q variant, this density could accommodate the side chains of either histidine or glutamine, but in a different conformation to that previously assigned to H50 in the wild-type structure. The presence of both H50 and Q50 residues in the α-synuclein filaments was also shown by mass spectrometry (Fig. S3). Analysis of α-synuclein filaments from the frontal cortex of the H50Q case showed a filament structure consistent with that from the amygdala, as judged by 2D class averaging (Fig. S5B).

**Fig. 3:**
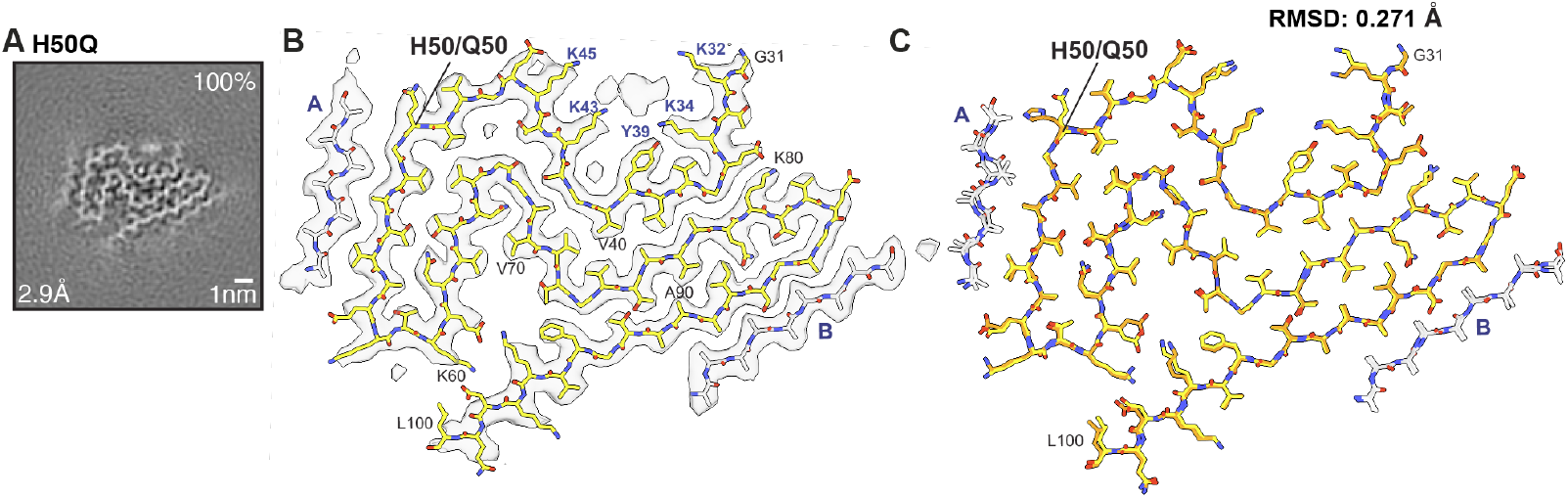
Cryo-EM structure of H50Q α-synuclein filaments from human brain. **(A)** Cross-section through the cryo-EM reconstruction, perpendicular to the helical axis and with a projected thickness of approximately one rung, for the α-synuclein singlet from a case with *SNCA* variant H50Q. Scale bar, 1 nm. **(B)** Cryo-EM density map (transparent grey) and atomic model of the singlet filaments. The model for α-synuclein is shown in yellow and that of islands A and B is in grey. **(C)** Overlay of the atomic models of the α-synuclein singlet from this case (yellow) and the Lewy fold singlet of idiopathic PD (orange). Density islands are in grey.

### Structures of α-synuclein filaments from homozygous mice transgenic for human A53T α-synuclein (line M83)

Last, we determined the cryo-EM structures of α-synuclein filaments from pooled brainstems and spinal cords of 45- and 47-week-old homozygous M83 transgenic mice, which express human mutant A53T α-synuclein down-stream of the murine prion protein promoter [29] (Fig. 4 and fig. S7). These mice suffered from weight loss and severe hindlimb paralysis. Most filaments were twisted by cryo-EM 2D class averaging and made of a single protofilament, with crossover distances of ∼2,000 Å; a minority of doublets with crossover distances of ∼1,100 Å was also observed. Structure determination of the singlets and doublets to resolutions of 2.7 Å and 3.0 Å, respectively, revealed left-handed helical twists and a shared protofilament fold, unlike any previously reported structures of α-synuclein filaments. We named it the mouse A53T (mA53T) fold.

**Fig. 4:**
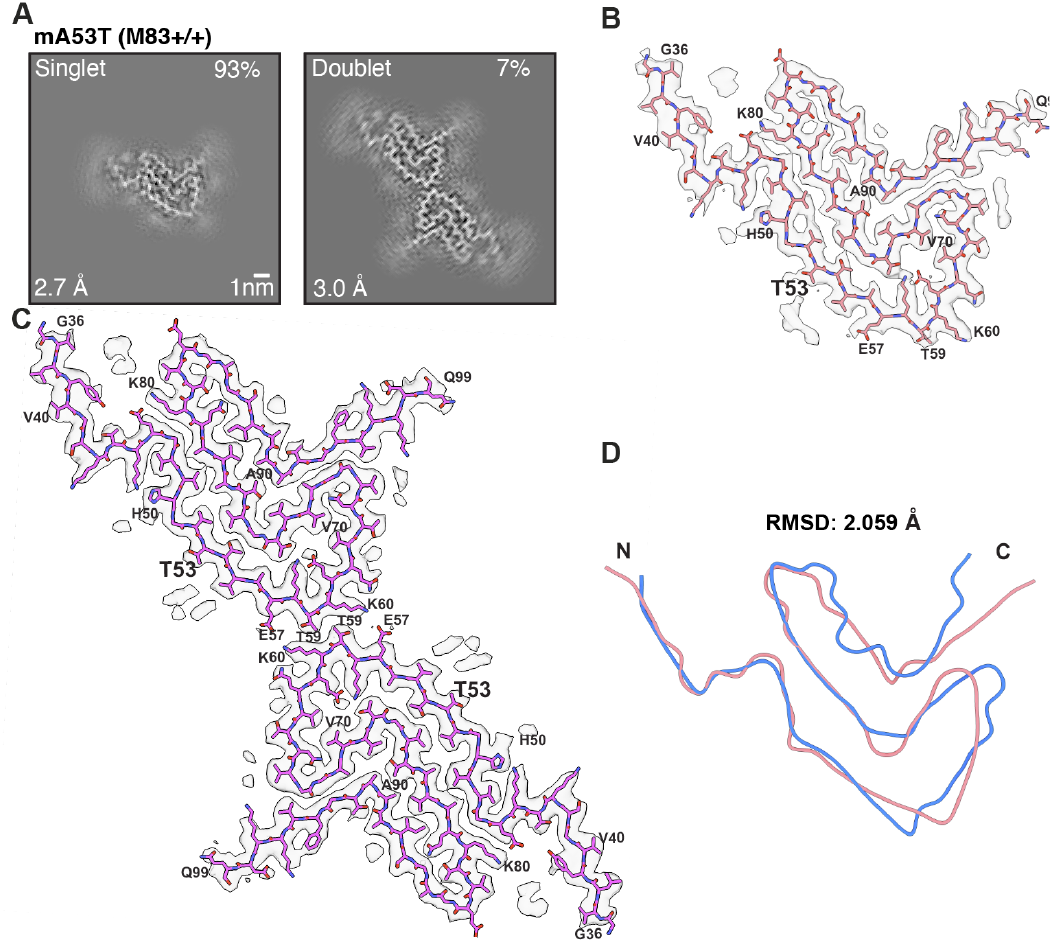
Cryo-EM structures of A53T α-synuclein filaments from M83 mice. **(A)** Cross-sections through the cryo-EM reconstructions, perpendicular the helical axis and with a projected thickness of approximately one rung, are shown for α-synuclein singlet and doublet filaments extracted from homozygous M83 mice. Scale bar, 1 nm. **(B)** Cryo-EM density map (transparent grey) and atomic model of the singlet filaments (magenta). **(C)** As in B, but for the doublet, with the atomic model in pink. **(D)** Back-bone superposition of the mA53T singlet (pink) and chain H protofilament from an atypical case of MSA (PDB:9OBP) [33] (blue).

The protofilament core of the mA53T fold extended from residues G36-Q99, in the absence of an island. It comprised eight β-strands that extended from residues 39-42, 48-50, 52-58, 60-66, 69-71, 74-80, 87-90 and 94-98 of α-synuclein. Additional non-proteinaceous densities were present in the pocket formed by residues G36-E46 and K80-E83, which may correspond to cofactors and/or post-translational modifications. In the mA53T doublets, protofilaments were related by 2-start helical symmetry and the dimer interface was formed by residues E57, T59 and K60 from each protofilament. The doublets were stabilised by two pairs of salt bridges between residues E57 and K60 of α-synuclein.

The mA53T protofilament fold differed from the Lewy fold of idiopathic PD, PDD and DLB with an APD of 76% and a backbone RMSD of 15.135 Å (Fig. S2D). Of known human brain-extracted α-synuclein folds it most closely resembled one protofilament from an atypical case of MSA [33], with an APD of 38% and a backbone RMSD of 2.059 Å (Fig. S2E). The mA53T protofilament fold differed from the folds of A53T protofilaments assembled *in vitro* and seeded in heterozygous M83 mice [37], but it shared the dimerisation interface with that of an *in vitro* assembled recombinant A53T α-filament (PDB:6LRQ) [27]. Unlike the positively charged pockets with unidentified non-proteinaceous densities in human brain-extracted filaments, the pocket with an unidentified density in the mA53T fold was essentially charge neutral and similar to the N-pocket in structures of recombinant α-synuclein polymorph 1a, where thioflavin T and Pittsburgh compound B have been shown to bind [38].

The mA53T fold can readily accommodate mouse α-synuclein that has a threonine at position 53 and differs from the core sequence only at the exposed site S87N, consistent with the presence of mouse α-synuclein in the mA53T fold, as determined by mass spectrometry (Fig. S3). Likewise, with threonine 53 pointing to the solvent, the mA53T fold can also accommodate wildtype human α-synuclein. The presence of mouse α-synuclein in the mA53T fold sets it apart from the fold of seeded filaments in heterozygous M83 mice (PDB 9RZF) [37], in which the side chain of S87 was buried inside a tight turn with little room for the larger side chain of N87 of mouse α-synuclein.

## Discussion

Contrary to expectations from studies with recombinant α-synuclein [24-28], filaments with mutations A53T and G51D from human brains adopt the Lewy fold that was previously described in idiopathic PD, PDD and DLB [23]. However, they lack density for island A, have a left-handed twist and form dimers (Fig. 5). These are the first structures of α-synuclein filaments from the brains of individuals with inherited PD. By contrast, filaments of α-synuclein with variant H50Q were like those from idiopathic PD; they comprised a single protofilament with a right-handed twist and a density for island A.

**Fig. 5:**
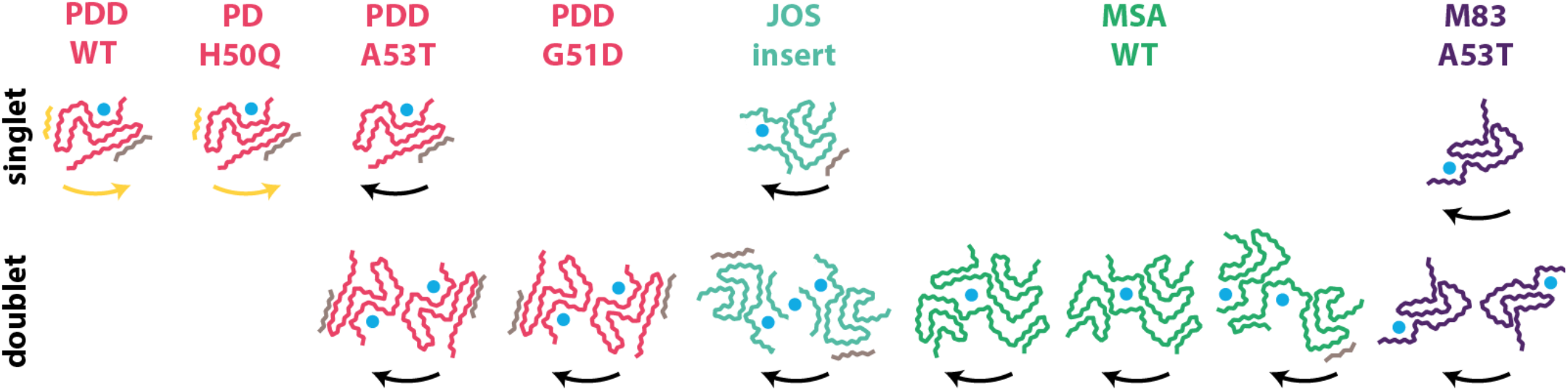
Comparison of the structures of brain-derived α-synuclein filaments. Filaments with the Lewy fold are shown in pink, filaments with the JOS fold in teal, filaments with the MSA folds (typical and atypical) in green and filaments with the mA53T fold (homozygous M83 mice) in purple. The density of island A is shown in yellow; densities of other islands are shown in grey. Unidentified cofactor densities are indicated with cyan circles. The handedness of the filaments is indicated with arrows. A density for island A correlates with the presence of a right-handed helical twist.

What do these structures tell us about the pathogenicity of *SNCA* mutations? The finding that H50Q α-synuclein filaments are like those from idiopathic PD supports the conclusion that this variant is not pathogenic [22]. Therefore, the individual with this amino acid change may have suffered from idiopathic PD, in agreement with the absence of a family history of PD. This observation contrasts with those for mutations A53T and G51D, where all carriers develop PD and family members without these mutations do not develop PD.

The structures of α-synuclein filaments with mutations G51D and A53T from human brains suggest a direct link between protofilament dimerisation, a left-handed helical twist and dominantly inherited PD. Dimers of α-synuclein filaments with a left-handed twist are also present in MSA (typical and atypical) [30-33] and JOS [34] (Fig. 5). Through avidity and a doubling of the number of hydrogen bonds along the helical axis, filaments made of two protofilaments are probably more stable than those with a single protofilament. Moreover, most filaments assembled from recombinant α-synuclein also adopt doublets with a left-handed twist [39]. Increased stability of α-synuclein filaments may contribute to their persistence in brain cells and hence cause neurodegeneration and disease. Dimerisation of the left-handed Lewy fold may also be necessary for positive Gallyas-Braak silver staining. Thus, brain sections from the A53T and G51D cases were silver-positive, whereas sections from the H50Q case were silver-negative. The Lewy pathology of idiopathic PD is silver-negative, whereas the α-synuclein inclusions of JOS and MSA are Gallyas-Braak silver-positive [40,41].

In the Lewy fold of idiopathic PD, PDD and DLB, both alanine 53 and glycine 51 form part of the interface with peptide-like density island A. The larger side chains of threonine and aspartate in cases with mutations A53T and G51D will disrupt this tightly packed interface, in agreement with the observation that the density of island A is absent from the singlets with mutation A53T. A weak density in its place can be explained by the predominantly non-polar nature of the interacting surface patch, which is preserved by the A53T mutation. This patch may interact with hydrophobic residues in the fuzzy coat of the α-synuclein fold and/or with other molecules. In contrast, the introduction of a negative charge in the G51D mutation will abolish hydrophobic interactions with this patch. The presence of island A will prevent protofilament dimerisation because of steric clashes with a second protofilament. Thereby, mutations A53T and G51D may lead to filament dimerisation through disruption of the interactions with the molecules responsible for the density of island A. The observation that the left-handed A53T singlet filament has a rearrangement in the Z-position of its backbone in the region opposite island A relative to the right-handed wild-type Lewy fold suggests that the interaction with island A may also affect the handedness of the filaments.

Disease-causing missense mutations in *SNCA* other than A53T and G51D map to either the exterior of the Lewy fold near the cofactor-binding site (A30G, A30P, E46K) [3,4,10] or the interface with island A (A53E, A53V) [8,9]. Like A53T, mutations A53E and A53V probably lead to the displacement of island A, resulting in dimerisation of the Lewy fold. The molecular identity of island A remains unknown. If some α-synuclein protein chains contribute to both the filament core and island A, with their linker going over the cofactor binding site and the dimerisation interface, it will enforce the incompatibility between the presence of island A and dimerisation of the Lewy fold protofilament. It provides a rationale for how mutations A30G, A30P and E46K might affect linker interactions. The α-synuclein filaments with variants K58N [42], T72M [43] and E83Q [44] are also compatible with the Lewy fold; K58N and E83Q map to the exterior of the Lewy fold and T72M maps to a large cavity inside that fold. Similarly, α-synuclein variants G14R [45] and V15A [46] are compatible with the Lewy fold; they are in the potential linker between island A and the core of the Lewy fold. It follows that α-synuclein filaments from all known PD-causing mutations may adopt the Lewy fold or variants thereof.

Even though the left-handed mA53T fold of α-synuclein filaments extracted from the brains of homozygous M83 mice is unlike any known human α-synuclein fold, it is more closely related to MSA protofilament folds than to the Lewy fold. It follows that transgenic mouse line M83 is a better model for MSA than for PD, despite the fact that it is transgenic for human mutant A53T α-synuclein, which causes PDD in humans.

This may explain why the intracerebral injection of MSA seeds into heterozygous M83 mice led to more neuronal α-synuclein inclusions and a more aggressive form of neurological disease than the injection of PD seeds [47-49]. Unlike their homozygous counterparts, heterozygous M83 mice do not develop inclusions or symptoms unless challenged with α-synuclein seeds. The intracerebral injection of recombinant wildtype α-synuclein filaments into heterozygous M83 mice produced some oligodendroglial α-synuclein inclusions reminiscent of MSA, but the structures of the recombinant filaments were distinct from those of MSA [30,37]. Moreover, the seeded filaments only partially retained the structures of the seeds and the structures of seeded α-synuclein filaments differed from those of filaments from homozygous M83 mice. It remains to be determined if MSA seeds can behave like true prions in this system by giving rise to seeded aggregates with the same structures as those of the seeds. The clinicopathological effects of the injection of human brain G51D α-synuclein seeds into heterozygous M83 mice resembled those of injected seeds from cases of idiopathic PD [50], consistent with our findings that G51D α-synuclein filaments are made of doublets of the Lewy fold.

## Conclusion

We establish that dimerisation of α-synuclein protofilaments with a left-handed helical twist leads to disease in cases of familial PD and dementia. This may be the gain-of-toxic function mechanism by which dominantly inherited *SNCA* mutations cause PD. We also underscore the need to develop methods by which to form α-synuclein filaments with the same structures as those from human brains. Even though limited progress towards this goal has been made for MSA, as illustrated by our structures of α-synuclein filaments from homozygous M83 mice and those by others from seeded heterozygous M83 mice [37], none of the existing methods come close to forming the Lewy fold of α-synuclein. Such methods will be crucial for elucidating the molecular mechanisms that underlie the formation of α-synuclein filaments with the Lewy fold and for the development of mechanism-based therapies for PD, PDD and DLB.

## Acknowledgements

We thank the patients and their families for donating brain tissues, T. Darling, I. Clayson and J. Grimmett for help with high-performance computing and the EM facility of the Medical Research Council (MRC) Laboratory of Molecular Biology for help with cryo-EM data acquisition. We also thank the advanced cryo-EM suite at the Van Andel Research Institute and the cryo-EM core facility at Michigan State University. We are grateful to S. Subramanian, X. Meng, G. Zhao and I. Ratnayake for their assistance with cryo-EM data acquisition, and the Van Andel Institute high-performance computing team for their support. We thank K. Cox and M. Jacobsen for assistance with silver staining. This work was supported by the MRC, as part of UK Research and Innovation (MC_UP_A025_1013 to S.H.W.S. and MC_1051284291 to M.G.). It was also supported by the Van Andel Institute and a West Michigan Neurodegenerative Diseases Program Pilot Award (MiP.017) to Y.Y.

## Author contributions

P.W.C., A.O., R.R., K.N., D.T.,D.K., H.W., H.H., B.G., N.H. and Z.J. identified patients, performed neuropathology and carried out genetic analysis; J.A.M. organised breeding and characterised mouse tissues; J.A.M., S.P.C, C.F. and Y.Y. prepared filaments and performed mass spectrometry; H.Z. and Y.Y. performed cryo-EM data acquisition; H.Z., T.V.H., A.G.M., Y.Y., and S.H.W.S. performed cryo-EM structure determination and analysis; Y.Y., S.H.W.S. and M.G. supervised the project; all authors contributed to writing the manuscript.

## Copyright statement

For the purpose of open access, the MRC Laboratory of Molecular Biology has applied a CC BY public copyright licence to any Author Accepted Manuscript version arising Data availability.

## Data availability

Cryo-EM maps were deposited to the EM Data Bank under accession codes EMD-76266 for A53T-singlet, EMD-76267 for A53T-doublet, EMD-70252 for G51D (case 1), EMD-70253 for H50Q, EMD-76268 for mAT53-singlet and EMD-76269 for mAT53-doublet. Corresponding refined atomic models were deposited to the PDB under accession codes 12AP for A53T-singlet, 12AQ for A53T-doublet, 9O9J for G51D (case 1), 9O9K for H50Q, 12AR for mA53T-singlet and 12AT for mA53T-doublet. Please address requests for materials to Y.Y., S.H.W.S. and M.G.

## Materials and Methods

### Genetics

DNA was extracted from blood of the A53T case [34], frontal cortex of G51D cases 1 and 2 [20] and amygdala of the H50Q case. For the A53T case, targeted gene panel sequencing was used to screen for genes associated with familial PD and other neurodegenerative diseases using the Torrent platform (Thermo Fisher Scientific). The variant in *SNCA* that encodes A53T α-synuclein was confirmed by Sanger sequencing. Copy number variation in *SNCA* was evaluated using multiplex ligation-dependent probe amplification. For the G51D and H50Q cases, DNA was sequenced using Illumina short-read whole genome sequencing, with a mean coverage of 30x. DNA from each sample was also genotyped with the Illumina Infinium Global Diversity Single Nucleotide Polymorphism Array with custom content for neurodegenerative diseases (Neurobooster) [51].

### Neuropathology

Human brains were examined neuropathologically following standard Queen Square Brain Bank protocols. Eight μm thick sections were cut from formalin-fixed, paraffin-embedded tissue blocks sampled from representative brain regions. Immunohistochemical staining for α-synuclein (MA1-90342, Thermo Fisher Scientific, 1:500), amyloid-β (M0872, Dako, Abnova, 1:500) and phosphorylated tau (MN1020, Thermo Fisher Scientific, 1:600) was performed on a Menarini automated staining platform following the manufacturers’ guidelines. Biotinylated secondary antibodies were used with horseradish-conjugated streptavidin complex and diaminobenzidine as the chromogen. To visualise inclusions, sections from the cerebral cortex were also impregnated according to Gallyas-Braak [52,53]. All sections were counterstained with haematoxylin.

### Transgenic mice

The M83 transgenic mouse line, which expresses human mutant A53T α-synuclein under the control of the murine prion protein promoter on a mixed C57BL6/C3H background [29], was purchased from the Jackson Laboratory (stock number 004479). Brainstems and spinal cords of symptomatic homozygous mice were used for cryo-EM and mass spectrometry. All experiments were carried out in compliance with the Animals (Scientific Procedure) Act of 1986 and were approved by the local Animal Welfare and Ethical Review Board.

### Filament extraction

Sarkosyl-insoluble material was extracted from temporal cortex (case of A53T α-synuclein), frontal cortex (cases of G51D and H50Q α-synuclein), amygdala (case of H50Q α-synuclein) and pooled brainstems and spinal cords (homozygous M83 mice), essentially as described [54]. Briefly, tissues were homogenised in 20 vol (w/v) extraction buffer consisting of 10 mM Tris-HCl, pH 7.4, 0.8 M NaCl, 10-20% sucrose and 1 mM EGTA. Homogenates were brought to 2% sarkosyl and incubated for 60 min at 37° C. Following a 10 min centrifugation at 10,000 g, the supernatants were spun at 100,000 g for 60 min. The final pellets were resuspended in 100 μl/g of 20 mM Tris-HCl, pH 7.4, 50 mM NaCl.

### Mass spectrometry

Mass spectrometry was performed as described [55]. Sarkosyl-insoluble pellets were resupended in 200 ml hexafluoroisopropanol. Following a 3 min sonication at 50% amplitude (QSonica), they were incubated at 37° C for 2 h and centrifuged at 3,000 g for 5 min; the supernatants were transferred to new tubes before being dried by vacuum centrifugation. They were resuspended in 4 M urea and 50 mM ammonium carbonate (ambic), before being reduced with 5 mM dithiothreitol at 37° C for 30 min and alkylated in the dark at room temperature for 30 min with 10 mM iodoacetamide. LysC (Promega) was then added to the samples for 2h at 25° C. They were diluted to 1.5 M urea with 50 mM ambic and incubated with trypsin (Promega) at 30° C overnight. Digestion was stopped by the addition of formic acid to 0.5%, followed by centrifugation at 16,000 g for 5 min. The supernatants were desalted using C18 stage tips (3M Empore) packed with Poros oligo R3 (Thermo Fisher Scientific) that was produced in-house. Bound peptides were eluted stepwise with 30, 50 and 80% acetonitrile in 0.5% formic acid and partially dried in a Speed Vac (Savant). The peptide mixtures were analysed by LC-MS/MS using a Q Exactive Plus hybrid quadrupole Orbitrap mass spectrometer, as described [56].

### Cryo-EM image acquisition

Holey carbon grids (Quantifoil auR1.2/1.3, 300 mesh) were glow-discharged with an Edwards (S150B) sputter coater at 30 mA for 30 s. Three μl aliquots were applied to the grids and blotted for 3-5 s with filter paper at 100% humidity and 4° C using a Thermo Fisher Vitrobot Mark IV. Datasets of the A53T case, G51D case 1 and the H50Q case were acquired on a Thermo Fisher G4 microscope, equipped with a Falcon-4i detector and a Selectris-X energy filter (Thermo Fisher Scientific) with a slit width of 10 eV to remove inelastically scattered electrons. The G51D case 2 dataset was collected on a Titan Krios G3 microscope with a Gatan K3 detector in super-resolution counting mode, using a Bioquantum energy filter with a slid width of 20 eV. The M83 mouse dataset was collected on a Titan Krios G2 microscope with a Gatan K3 detector in super-resolution counting mode, using a BioQuantum energy filter with a slid width of 20 eV. Images were recorded with a dose of 40 electrons per Å^*2*^ (50 electrons per Å^*2*^ for mA53T)

### Cryo-EM image processing

All super-resolution frames were gain-corrected, binned by a factor of 2, aligned, dose-weighted and then summed into a single micrograph using RELION’s own implementation of MotionCor2 [57]. Contrast transfer function (CTF) parameters were estimated using CTFFIND-4.1 [58]. Subsequent image-processing steps were performed using helical reconstruction methods in RELION [59,60]. Filaments were picked manually. Reference-free 2D classification was performed to identify homogeneous segments for further processing. Initial 3D reference models were reconstructed *de novo* from 2D class averages [61] using an estimated rise of 4.75 Å and helical twists according to the observed crossover distances of filaments in the micrographs. 3D classification was used to select optimal particles for each structure. To increase the resolution of the reconstructions, Bayesian polishing and CTF refinement were performed [62]. Blush regularisation was applied to improve the quality of the reconstructions [63]. Final 3D reconstructions, after auto-refinement, were sharpened using the standard post-processing procedures in RELION, and resolutions were calculated from Fourier shell correlations at 0.143 between the two independently refines half-maps, using phase-randomisation to correct for convolution effects of a generous, soft-edged solvent mask. Further details of the data acquisition and processing are given in the Table S1.

### Atomic model building

Atomic models were built in Coot [64] using the shared substructures of the Lewy fold [23] (PDB: 8A9L) as the template. Coordinate refinements were performed using *Servalcat* [65]. Final models were obtained using refinement of only the asymmetric unit against the half-maps in *Servalcat*.

---

**Fig. S1:**
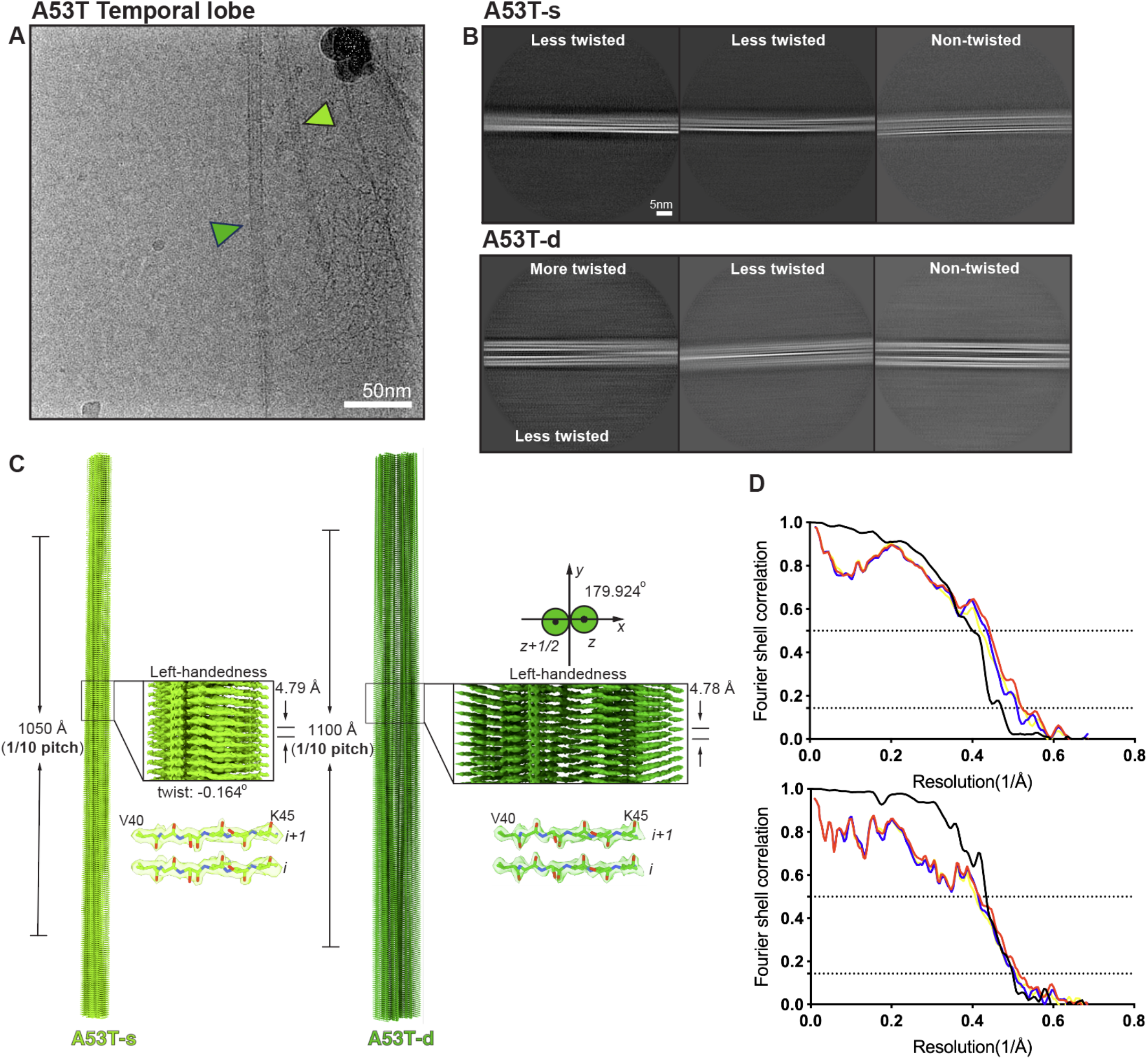
Cryo-EM of A53T α-synuclein filaments from human brain. **(A)** An example micrograph; a singlet is indicated with a light green arrow, a doublet with a dark green arrow. **(B)** 2D class averages of more twisted, less twisted and untwisted filaments for the singlet and the doublet; **(C)** Side views of the 3D reconstructions of the singlet (light green) and doublet (dark green) showing separation of the β-strands; **(D)** Fourier shell correlation (FSC) curves between two independently refined half maps (black), between the refined atomic model and the cryo-EM reconstruction (red), and between the two half maps (blue and yellow) of the singlet (top) and doublet (bottom).

**Fig. S2:**
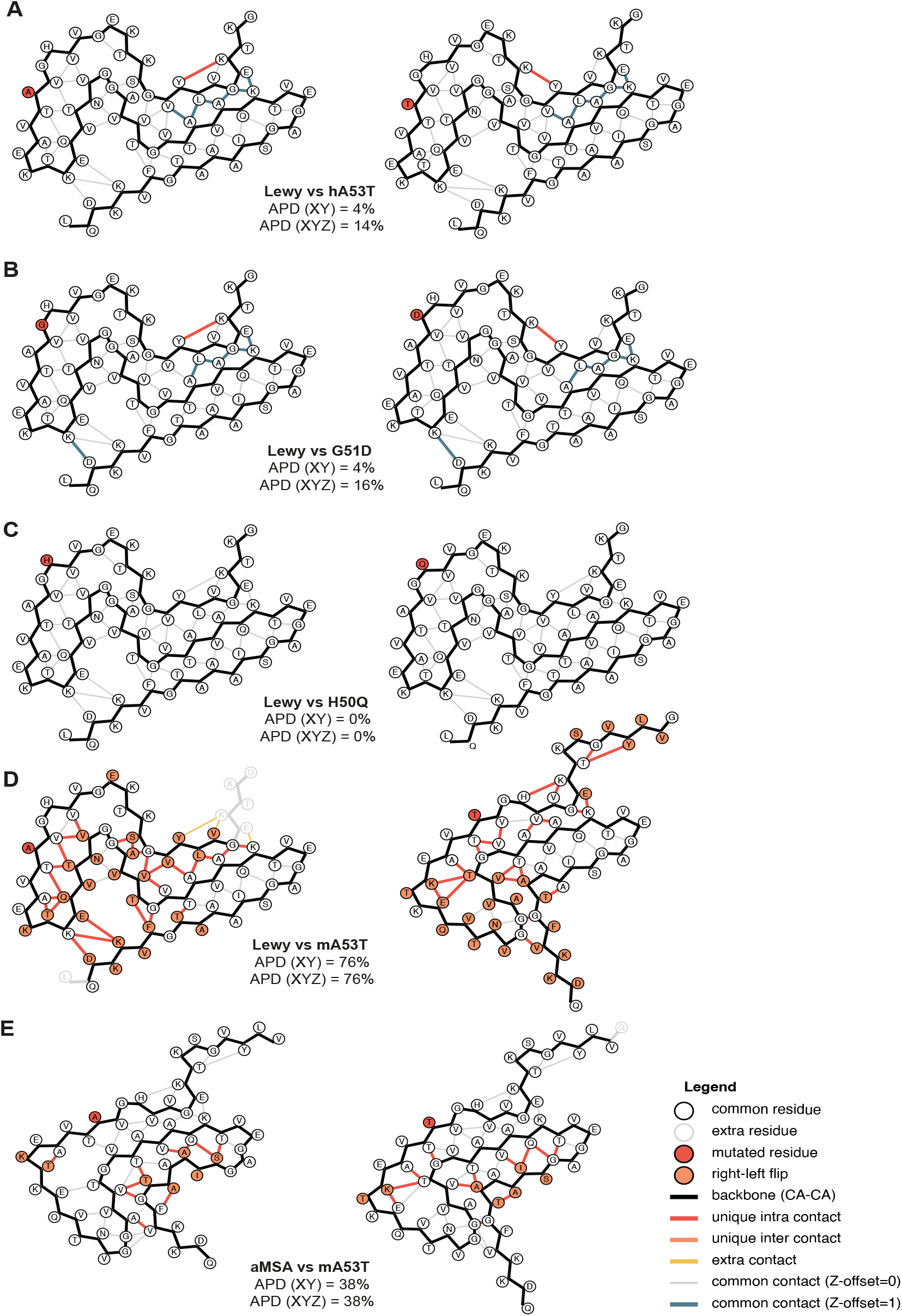
Amyloid packing differences. Amyloid packing differences (APDs) are shown between the Lewy fold from idiopathic PD, PDD and DLB and the protofilament folds for human A53T (hA53T) **(A)**, G51D **(B)**, H50Q **(C)** and murine A53T (mA53T) **(D)**, as well as between chain H protofilament from an atypical case of MSA (aMSA) and mA53T **(E)**.

**Fig. S3:**
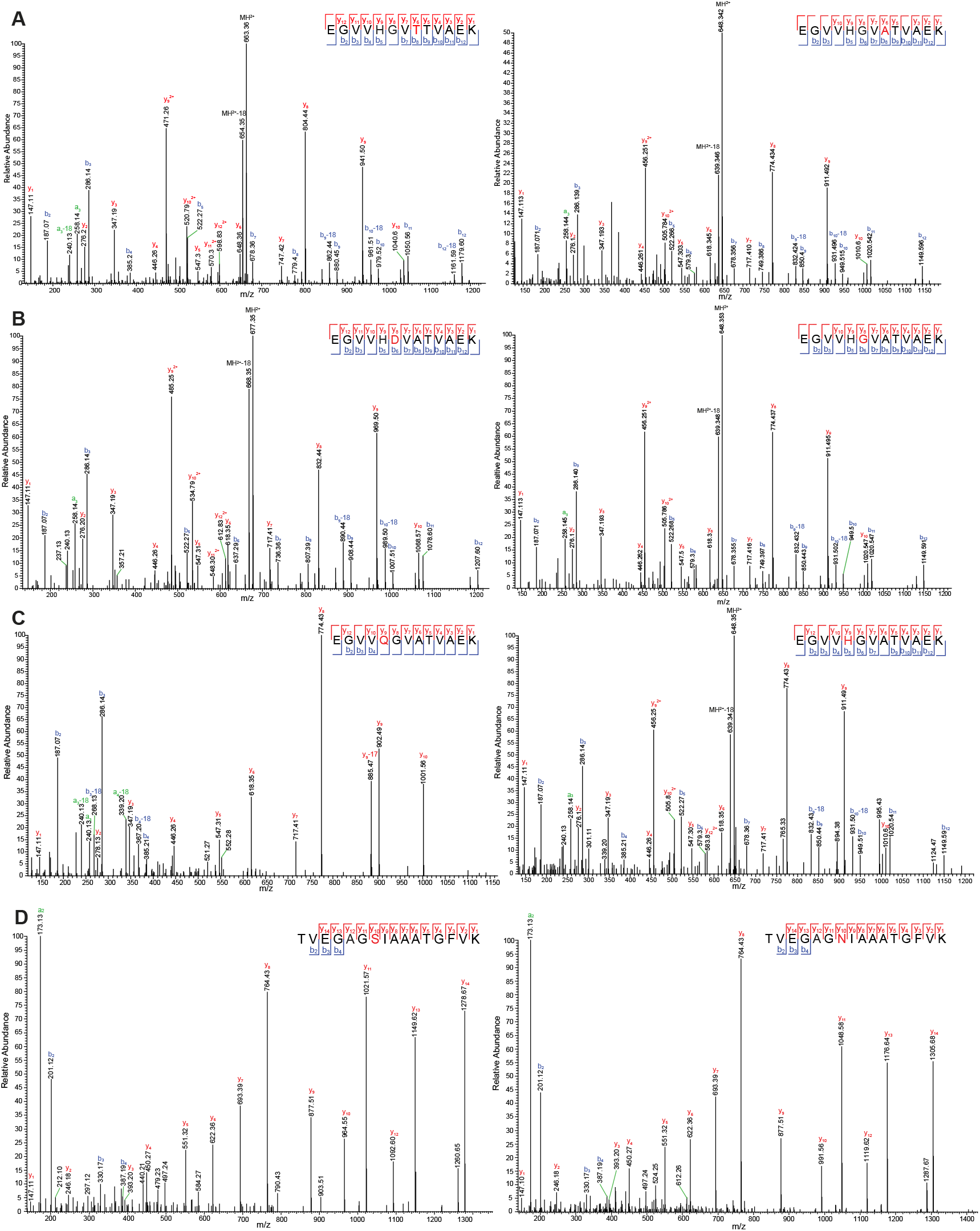
Mass spectrometry. Spectra of sarkosyl-insoluble fractions from the brains of individuals with *SNCA* mutations A53T **(A)** and G51D **(B)**, variant H50Q **(C)** and from mice homozygous for human mutant A53T α-synuclein (M83 mice) **(D)**. Peptide sequences differ by one amino acid (depicted in red) and are shown at the top of each panel with the corresponding higher-energy collisional dissociation (HCD) fragmentation spectra shown below; b ions (N-terminal; depicted in blue) and y ions (C-terminal; depicted in red) are generated during peptide fragmentation. The ions that have a neutral loss are shown in green.

**Fig. S4:**
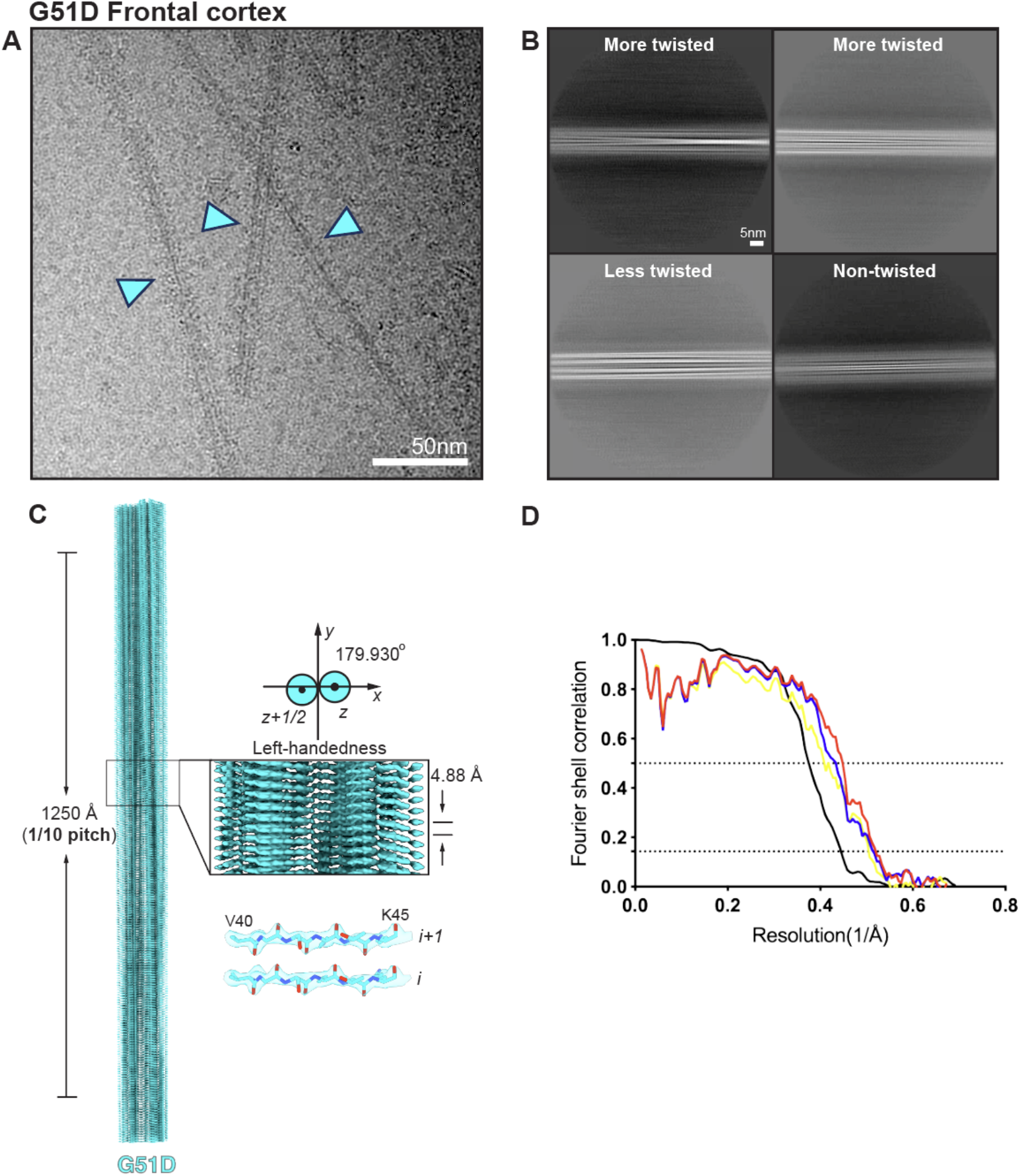
Cryo-EM of G51D α-synuclein filaments from human brain. **(A)** An example micrograph with G51D doublets indicated with cyan arrows; **(B)** 2D class averages of twisted and untwisted filaments; **(C)** Side view of the 3D reconstruction showing separation of the β-strands; **(D)** Fourier shell correlation (FSC) curves between two independently refined half maps (black), between the refined atomic model and the cryo-EM reconstruction (red), and between the two half maps (blue and yellow).

**Fig. S5:**
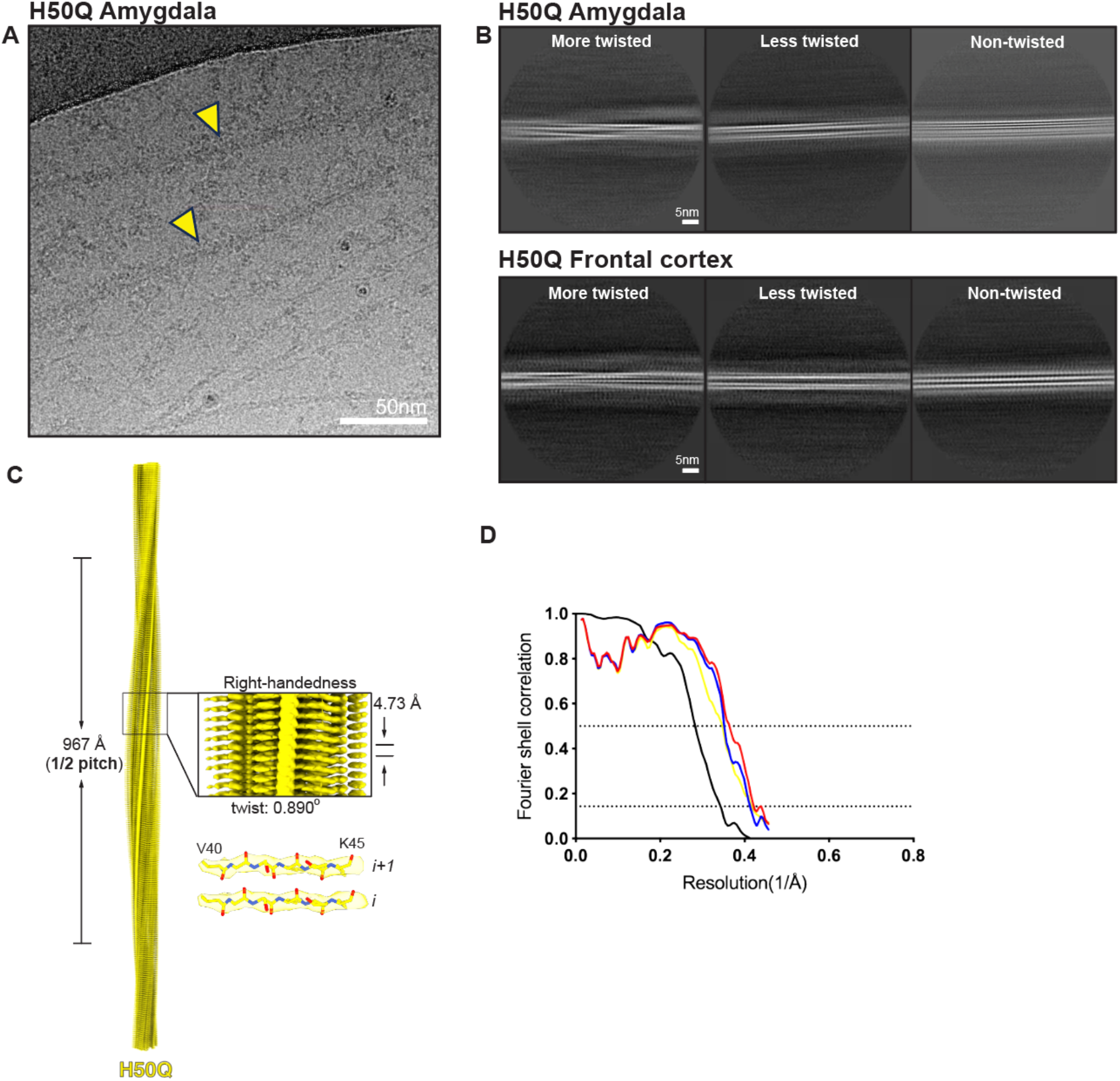
Cryo-EM of H50Q α-synuclein filaments from human brain. **(A)** An example micrograph, with H50Q singlets indicated with yellow arrows; **(B)** 2D class averages of twisted and untwisted filaments from amygdala and frontal cortex; **(C)** Side view of the 3D reconstruction showing separation of the β-strands; **(D)** Fourier shell correlation (FSC) curves between two independently refined half maps (black), between the refined atomic model and the cryo-EM reconstruction (red), and between the two half maps (blue and yellow).

**Fig. S6:**
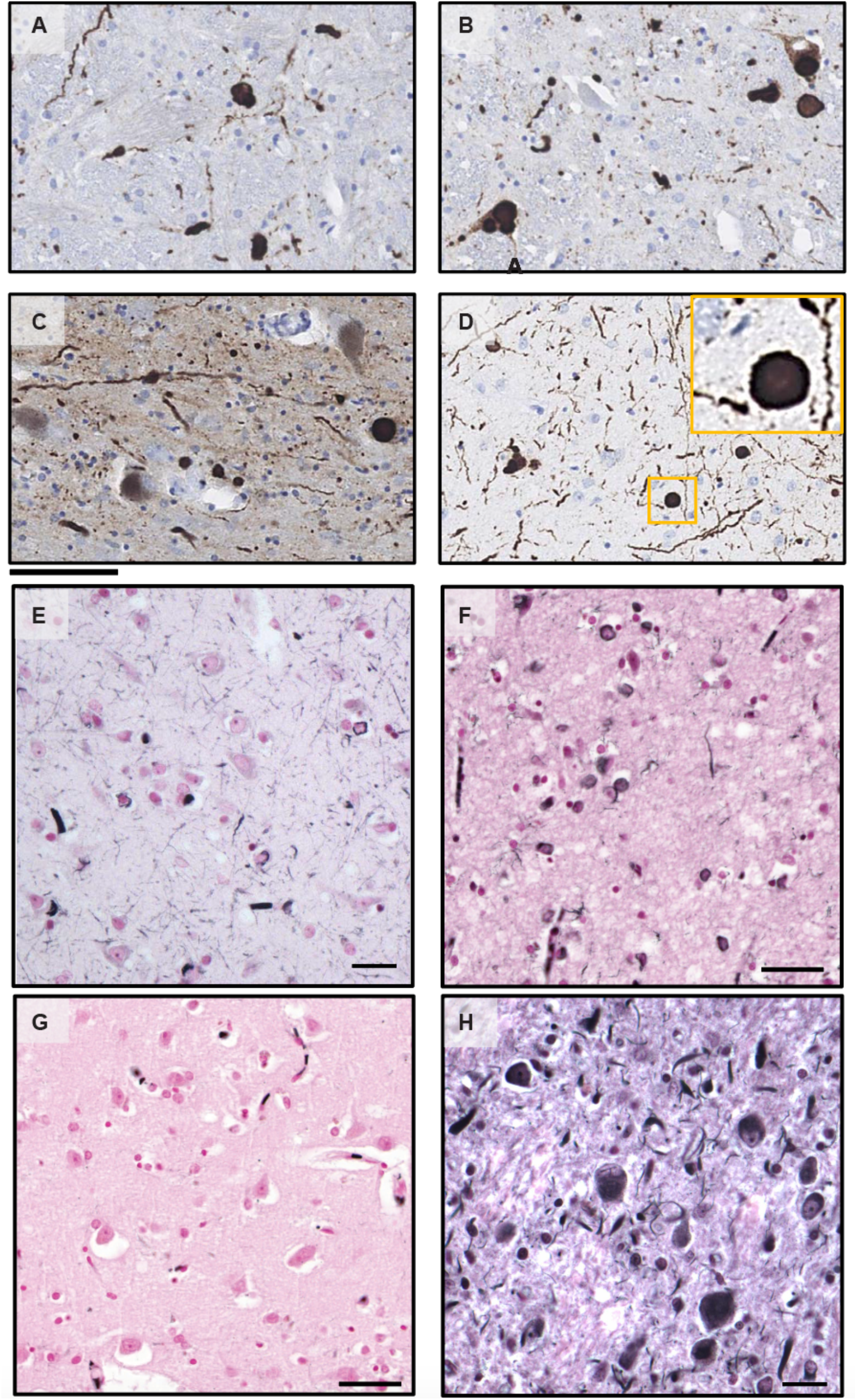
Immunostaining of α-synuclein inclusions from the H50Q case and Gallyas-Braak silver staining of A53T, G51D and H50Q cases. **(A-D)** Sections from brain regions contralateral to those used for cryo-EM structure determination were stained with anti-α-synuclein antibody MA1-90342 (1:500). There was widespread staining in nerve cells of the dorsal medulla (A), locus coeruleus (B), substantia nigra (C) and CA4 subregion of the hippocampus (D). The morphology of inclusions was that of typical Lewy bodies and Lewy neurites. **(E-H)** Gallyas-Braak silver staining of temporal cortex from the A53T case (E) and amygdala from G51D case 2 (F). No Gallyas-Braak staining of amygdala from the H50Q case (G) and strong staining of pons from a neuropathologically confirmed case of MSA (H). Scale bars: 80 μm (A-D); 50 μm (F); 25 μm (E,G,H); 20 μm (G).

**Fig. S7:**
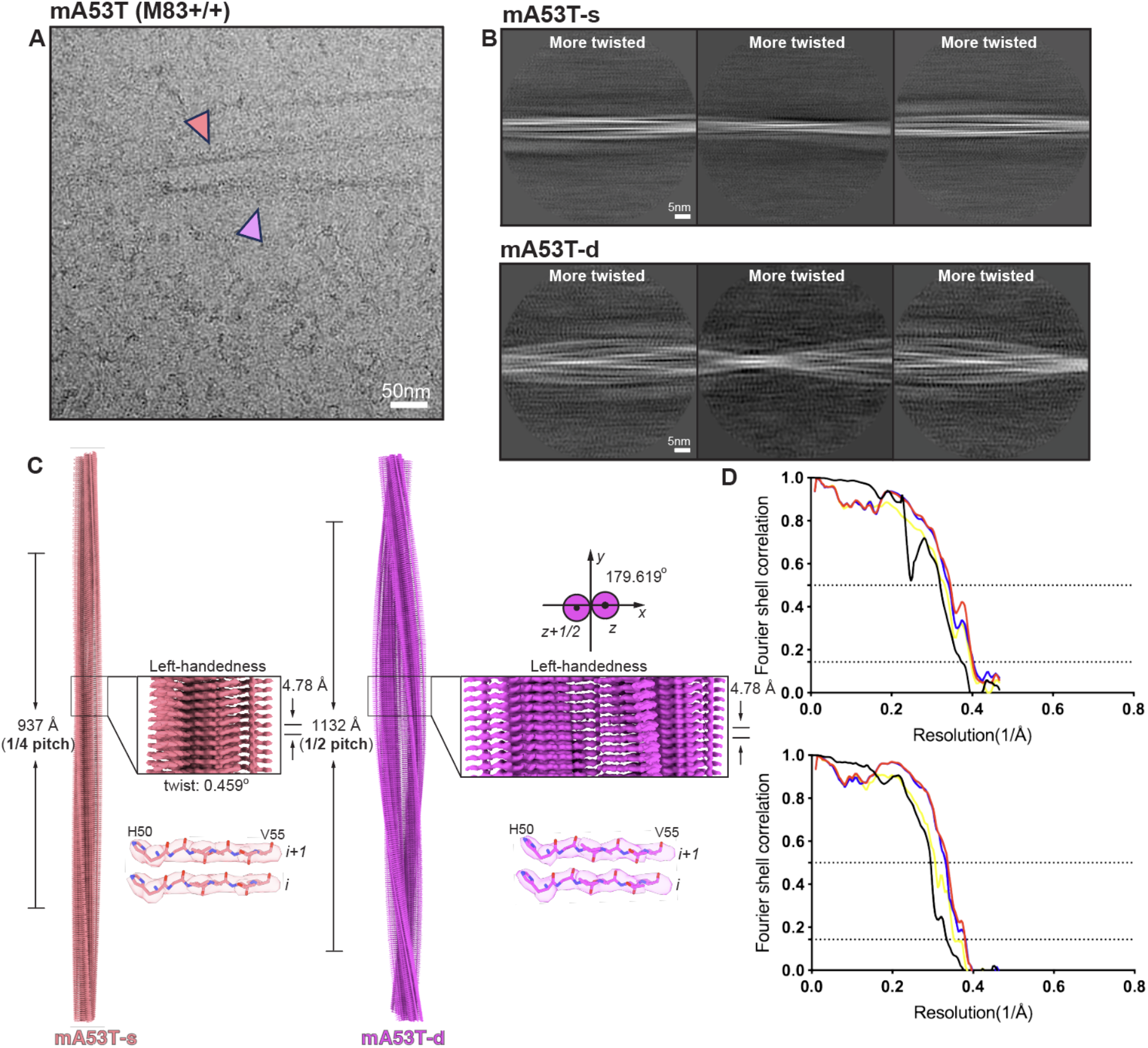
Cryo-EM of A53T α-synuclein filaments from M83 mice. **(A)** An example micrograph with singlets indicated with pink arrows and doublets indicated with purple arrows; **(B)** 2D class averages of twisted filaments; **(C)** Side view of the 3D reconstructions for the singlet (pink) and doublet (purple) protofilaments showing separation of the β-strands; **(D)** Fourier shell correlation (FSC) curves between two independently refined half maps (black), between the refined atomic model and the cryo-EM reconstruction (red), and between the two half maps (blue and yellow) for the singlet (top) and the doublet (bottom).

**Table S1:** Cryo-EM data acquisition and structure determination.

|  | **A53T-s** | **A53T-d** | **G51D (case1)** | **H50Q** | **mA53T-s** | **mA53T-d** |
| --- | --- | --- | --- | --- | --- | --- |
| EMDB | 76266 | 76267 | 70252 | 70253 | 76268 | 76269 |
| PDB | 12AP | 12AQ | 9O9J | 9O9K | 12AR | 12AT |
| **Data acquisition** |  |  |  |  |  |  |
| Electron gun | CFEG | CFEG | CFEG | CFEG | Krios G3 | Krios G3 |
| Detector | Falcon 4i | Falcon 4i | Falcon 4i | Falcon 4i | K3 | K3 |
| Energy filter slit (eV) | 10 | 10 | 10 | 10 | 20 | 20 |
| Magnification | 165,000 | 165,000 | 165,000 | 165,000 | 81,000 | 81,000 |
| Voltage (kV) | 300 | 300 | 300 | 300 | 300 | 300 |
| Electron dose ($e^-/\text{\AA}^2$) | 40 | 40 | 40 | 40 | 50 | 50 |
| Defocus range ($\mu\text{m}$) | 0.6 to 1.8 | 0.6 to 1.8 | 0.6 to 1.4 | 0.6 to 1.8 | 1.2 to 1.8 | 1.2 to 1.8 |
| Pixel size ($\text{\AA}$) | 0.731 | 0.731 | 0.744 | 0.724 | 1.07 | 1.07 |
| **Map refinement** |  |  |  |  |  |  |
| Symmetry imposed | C1 | C1 | C1 | C1 | C1 | C1 |
| Initial particle images (no.) | 144,525 | 197,560 | 362,549 | 618,647 | 230,923 | 16,643 |
| Final particle images (no.) | 39,772 | 180,726 | 247,047 | 200,126 | 59,954 | 10,056 |
| Map resolution ($\text{\AA}$) | 2.1 | 2.1 | 2.3 | 2.9 | 2.9 | 3.1 |
| FSC threshold | 0.143 | 0.143 | 0.143 | 0.143 | 0.143 | 0.143 |
| Helical twist ($^\circ$) | -0.164 | 179.924 | 179.93 | 0.89 | -0.46 | 179.62 |
| Helical rise ($\text{\AA}$) | 4.79 | 2.39 | 2.44 | 4.73 | 4.78 | 2.39 |
| **Model refinement** |  |  |  |  |  |  |
| Model resolution ($\text{\AA}$) | 2.1 | 2.1 | 2.3 | 2.9 | 2.7 | 3.0 |
| FSC threshold | 0.5 | 0.5 | 0.5 | 0.5 | 0.5 | 0.5 |
| Map sharpening $B$ factor ($\text{\AA}^2$) | -44.2 | -47.0 | -43.5 | -40.4 | -58.1 | -51.0 |
| **Model composition** |  |  |  |  |  |  |
| Non-hydrogen atoms | 2156 | 3282 | 4296 | 2835 | 1332 | 2220 |
| Protein residues | 316 | 474 | 632 | 430 | 192 | 320 |
| Ligands | 0 | 0 | 0 | 0 | 0 | 0 |
| **$B$ factors ($\text{\AA}^2$)** |  |  |  |  |  |  |
| Protein | 61.0 | 58.3 | 62.6 | 49.9 | 62.8 | 43.6 |
| **R.m.s. deviations** |  |  |  |  |  |  |
| Bond lengths ($\text{\AA}$) | 0.0087 | 0.0080 | 0.0092 | 0.0100 | 0.0096 | 0.0096 |
| Bond angles ($^\circ$) | 1.49 | 1.25 | 1.65 | 1.78 | 1.76 | 1.88 |
| **Validation** |  |  |  |  |  |  |
| MolProbity score | 1.81 | 1.57 | 0.99 | 2.26 | 1.85 | 1.60 |
| Clashscore | 5.93 | 2.89 | 1.37 | 7.75 | 6.61 | 5.07 |
| Poor rotamers (%) | 0 | 0 | 0.98 | 2.00 | 0 | 0 |
| **Ramachandran plot** |  |  |  |  |  |  |
| Favoured (%) | 92.00 | 92.00 | 97.33 | 87.50 | 91.94 | 95.16 |
| Allowed (%) | 8.00 | 8.00 | 2.67 | 12.50 | 8.06 | 4.84 |
| Disallowed (%) | 0 | 0 | 0 | 0 | 0 | 0 |

